# Structural and energetic profiling of SARS-CoV-2 antibody recognition and the impact of circulating variants

**DOI:** 10.1101/2021.03.21.436311

**Authors:** Rui Yin, Johnathan D. Guest, Ghazaleh Taherzadeh, Ragul Gowthaman, Ipsa Mittra, Jane Quackenbush, Brian G. Pierce

**Affiliations:** University of Maryland Institute for Bioscience and Biotechnology Research, Rockville, MD 20850, USA; Department of Cell Biology and Molecular Genetics, University of Maryland, College Park, MD 20742, USA

## Abstract

The SARS-CoV-2 pandemic highlights the need for a detailed molecular understanding of protective antibody responses. This is underscored by the emergence and spread of SARS-CoV-2 variants, including B.1.1.7, P1, and B.1.351, some of which appear to be less effectively targeted by current monoclonal antibodies and vaccines. Here we report a high resolution and comprehensive map of antibody recognition of the SARS-CoV-2 spike receptor binding domain (RBD), which is the target of most neutralizing antibodies, using computational structural analysis. With a dataset of nonredundant experimentally determined antibody-RBD structures, we classified antibodies by RBD residue binding determinants using unsupervised clustering. We also identified the energetic and conservation features of epitope residues and assessed the capacity of viral variant mutations to disrupt antibody recognition, revealing sets of antibodies predicted to effectively target recently described viral variants. This detailed structure-based reference of antibody RBD recognition signatures can inform therapeutic and vaccine design strategies.

## INTRODUCTION

Over the past year, the SARS-CoV-2 pandemic has resulted in a massive and growing global death toll and disease burden. A number of vaccines (Krammer, 2020), monoclonal antibodies (Jiang et al., 2020), and small molecule therapies (Simonis et al., 2021) that target SARS-CoV-2 have been developed. However, viral variants have raised the possibility of viral escape from, or reduced efficacy of, current vaccines and therapeutics (Liu et al., 2021a; Madhi et al., 2021; Starr et al., 2021; Wang et al., 2021b; Wang et al., 2021c; Wu et al., 2021).

Several recent studies have used in vitro experimental approaches to test human sera (Greaney et al., 2021a; Wang et al., 2021b) and sets of monoclonal antibodies (Greaney et al., 2021b; Liu et al., 2021b; Starr et al., 2021; Wang et al., 2021b) to profile SARS-CoV-2 antibody resistance. The rapidly expanding set of experimentally determined structures of antibodies targeting the spike glycoprotein provides the opportunity to use computational biology tools to map key features of antibody-spike recognition. At the same time, the impact of viral variability can be predicted, which can provide insights into effective targeting and neutralization of SARS-CoV-2 and enable selection and engineering of anti-spike therapeutics and vaccines.

Here we report detailed structural analysis of a large set of high resolution antibody-spike complexes that have been collected in our database, CoV3D (Gowthaman et al., 2021). Structure-based mapping of antibody footprints on the receptor binding domain (RBD) and unsupervised clustering led to the identification of four major antibody groups based on their recognition signatures. These antibody-spike complexes were assessed for key energetic features using computational alanine mutagenesis of all RBD interface residues to identify shared and distinct binding hotspots on the RBD. The structure-based antibody clusters were also assessed both for residue conservation with SARS-CoV-1, and predicted effects of individual RBD substitutions from circulating SARS-CoV-2 variants, showing substantial differences between groups of RBD-targeting antibodies. These structural features and clusters can serve as a reference for rational vaccine design and therapeutic efforts, and updated antibody cluster information based on this analysis is available to the community on the CoV3D site: https://cov3d.ibbr.umd.edu/antibody_classification.

## RESULTS

### Clustering of antibody-RBD interaction modes

To identify common recognition modes and key features of antibody recognition of the spike glycoprotein, we analyzed a set of high resolution structures of antibody-spike complexes from the CoV3D database (Gowthaman et al., 2021), which were originally obtained from the Protein Data Bank (Rose et al., 2011). We focused on the SARS-CoV-2 RBD, which is the primary target of neutralizing antibodies (Zost et al., 2020) and is the target of the vast majority of structurally characterized SARS-CoV-2 antibodies. Structures were filtered by resolution (< 4.0 Å) and nonredundancy, resulting in 70 antibody-RBD complex structures, representing different antibody formats (heavy-light antibody, nanobody) and a range of IGHV genes (**Table S1**). Notably, these complex structures include multiple therapeutic monoclonal antibodies that have been under clinical investigation: REGN10933 and REGN10987 (casirivimab/imdevimab; REGN-COV2) (Weinreich et al., 2021), LY-CoV555 (bamlanivimab) (Chen et al., 2021), and S309 which is the basis for VIR-7831 (GSK4182136; sotrovimab) (Tuccori et al., 2020).

To assess prevalent or shared binding modes in antibody-RBD recognition, pairwise root-mean-square-distances (RMSDs) between antibody heavy chain and nanobody chain orientations were calculated after superposition of RBD coordinates into a common reference frame, and the RMSDs were input to hierarchical clustering analysis (**Figure 1**). This analysis identified a set of 17 complexes with a common binding mode and shared heavy chain germline genes (IGHV3-53, IGHV3-66), a feature that has been noted in previous studies describing SARS-CoV-2 antibody-RBD complex structures (Barnes et al., 2020b; Yuan et al., 2021). Other sets of co-clustered antibodies within the 8 Å RMSD cutoff were limited to antibody pairs, with the exception of a set of five antibodies, of which three (2-15, Ab2-4, C121) share the IGHV1-2 heavy chain germline gene, suggestive of another germline-mediated binding mode. However, other antibodies possessing the IGHV1-2 germline gene exhibited distinct binding modes based on the clustering analysis (298, S2E12), indicating that the heavy chain CDR3 sequence and light chain are relevant factors for that orientation. An example of co-clustered antibodies based on this analysis is shown in **Figure 1B**, showing a shared RBD binding mode (heavy chain orientation RMSD: 2.9 Å) for neutralizing antibodies S304 (Piccoli et al., 2020) and EY6A (Zhou et al., 2020).

**Figure 1.**
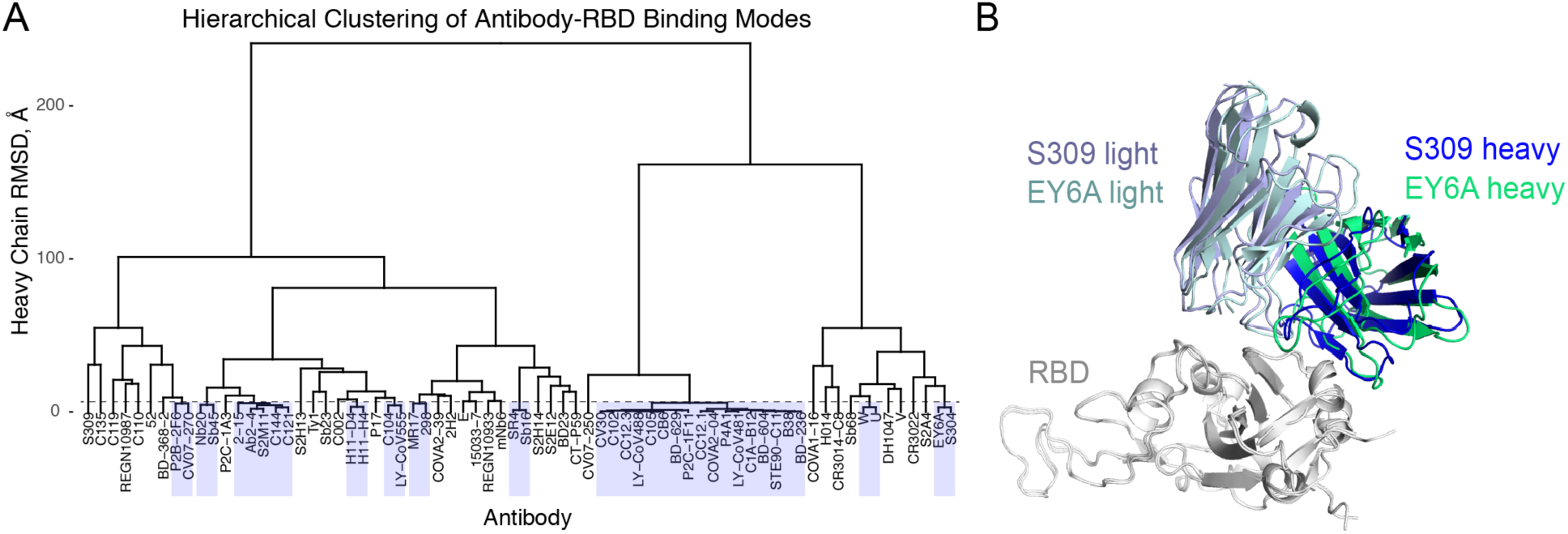
Hierarchical clustering of SARS-CoV-2 RBD antibody binding modes. (A) Pairwise root mean square distances (RMSDs) between heavy chain or nanobody binding orientations were determined for 70 antibody-RBD complex structures and used to perform hierarchical clustering. Boxes denote clusters containing multiple antibodies at distance cutoff of 7 Å (shown as dashed horizontal line). (B) Example of co-clustered antibodies S304 (PDB code 7JX3) (Piccoli et al., 2020) and EY6A (PDB code 6ZCZ) (Zhou et al., 2020) with a shared RBD binding mode (2.9 Å heavy chain orientation RMSD; far right cluster in panel (A)). Structures are superposed by RBD (gray), and S304 and EY6A heavy and light chains are colored separately as indicated.

### High resolution antibody footprinting and clustering analysis

To further delineate features underlying antibody-RBD recognition, we analyzed detailed antibody footprints on the RBD with unsupervised clustering, using the number of atomic contacts by an antibody to each RBD residue as input. Individual antibody footprints and resultant clusters are shown in **Figure 2**, along with calculated and previously reported properties of the antibodies for reference, including interface buried surface area (BSA), neutralization (SARS-CoV-2 neutralization or SARS-CoV-1/SARS-CoV-2 cross neutralization), ACE2 blocking, and capability to bind the RBD in the context of the closed (or down) spike conformation. This separated the antibodies into four main clusters; these are similar but not identical to previously described SARS-CoV-2 antibody classifications described by Barnes et al. (Barnes et al., 2020a), which are shown as the “BBclass” colored sidebar in **Figure 2**. Inspection of the heatmap indicates that Clusters 1 and 4 are most distinct, which is supported by high bootstrap confidence levels (100% and 99% respectively; **Figure S1**), while Clusters 2 and 3 are more diverse, and have bootstrap confidence levels of 87% and 83% (**Figure S1**). Visualization the distribution of the antibody positions on the RBD surface (**Figure 3**) shows that Clusters 1 and 2 are spatially proximal and overlap with the ACE2 binding site, and the relatively constrained positions of Cluster 1 antibodies are reflective of our RMSD-based analysis and known conserved binding mode of that set. Cluster 3 extends to the RBD hinge and N-glycan at RBD position N343, while Cluster 4 occupies a distinct region of the RBD. Principal component analysis using the antibody atom contact data as input enabled visualization of the antibody distributions along the first two principal components, which collectively represent approximately 50% of the data (**Figure S2**), and generally supports the hierarchical clustering.

**Figure 2.**
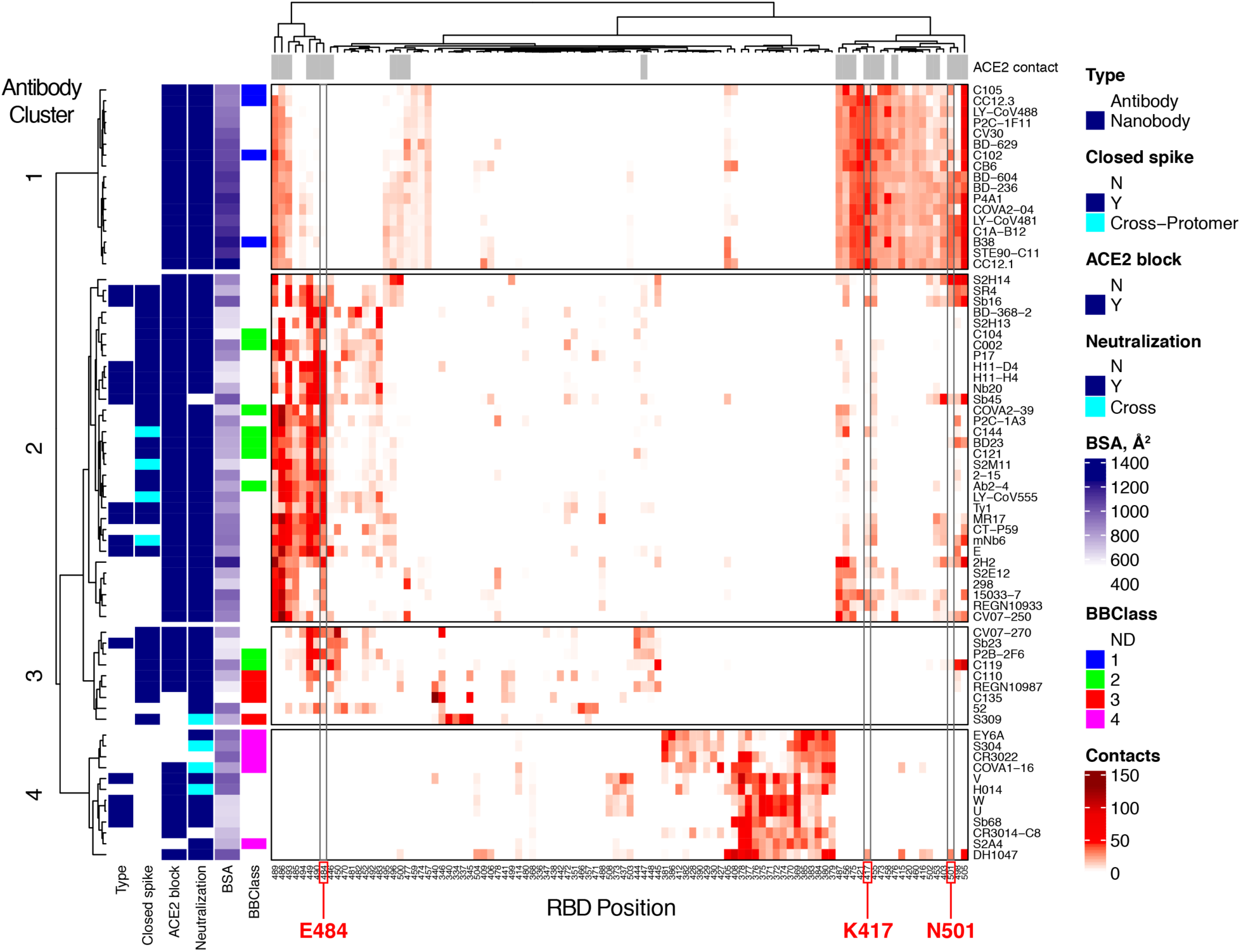
High resolution mapping and clustering of SARS-CoV-2 RBD antibody binding. RBD residue contact profiles were generated for each antibody based on number of antibody atomic contacts for each RBD residue within a 5 Å distance cutoff. RBD residues and antibodies are ordered using hierarchical clustering analysis, with dendrograms shown on top and left. The antibodies are separated into four major clusters based on contact profiles, and cluster numbers (1-4) are indicated on left. Contacts in heatmap are colored by number of RBD residue antibody atomic contacts, as indicated in the key. For reference, antibody type (Antibody: heavy-chain antibody, Nanobody: single-chain antibody), binding to RBD-closed spike conformation (Closed spike), ability to block ACE2 binding (ACE2 block), SARS-CoV-2 neutralization or SARS-CoV-2/SARS-CoV-1 cross-neutralization (“Y” and “Cross”, respectively, under Neutralization), interface buried surface area (BSA, Å^2^), and antibody classifications from a recent study (BBclass, with ND: antibody-RBD complex structure not described in the study)(Barnes et al., 2020a) are shown on the left sidebars. Closed spike binding and ACE2 blocking were calculated based on the structures, as described in the Methods. The top bar above the heatmap indicates RBD residues contacted by ACE2 (5 Å distance cutoff) in an ACE2-RBD complex structure (PDB code 6LZG) (Wang et al., 2020). For clarity, 100 RBD residues are shown in heatmap; a heatmap with the full set of 139 contacted RBD residues which was used to cluster the antibodies in this figure is shown in **Figure S1**. RBD residues that are mutated in SARS-CoV-2 variants of concern (K417, E484, N501) are labeled at bottom and highlighted with gray boxes in heatmap.

**Figure 3.**
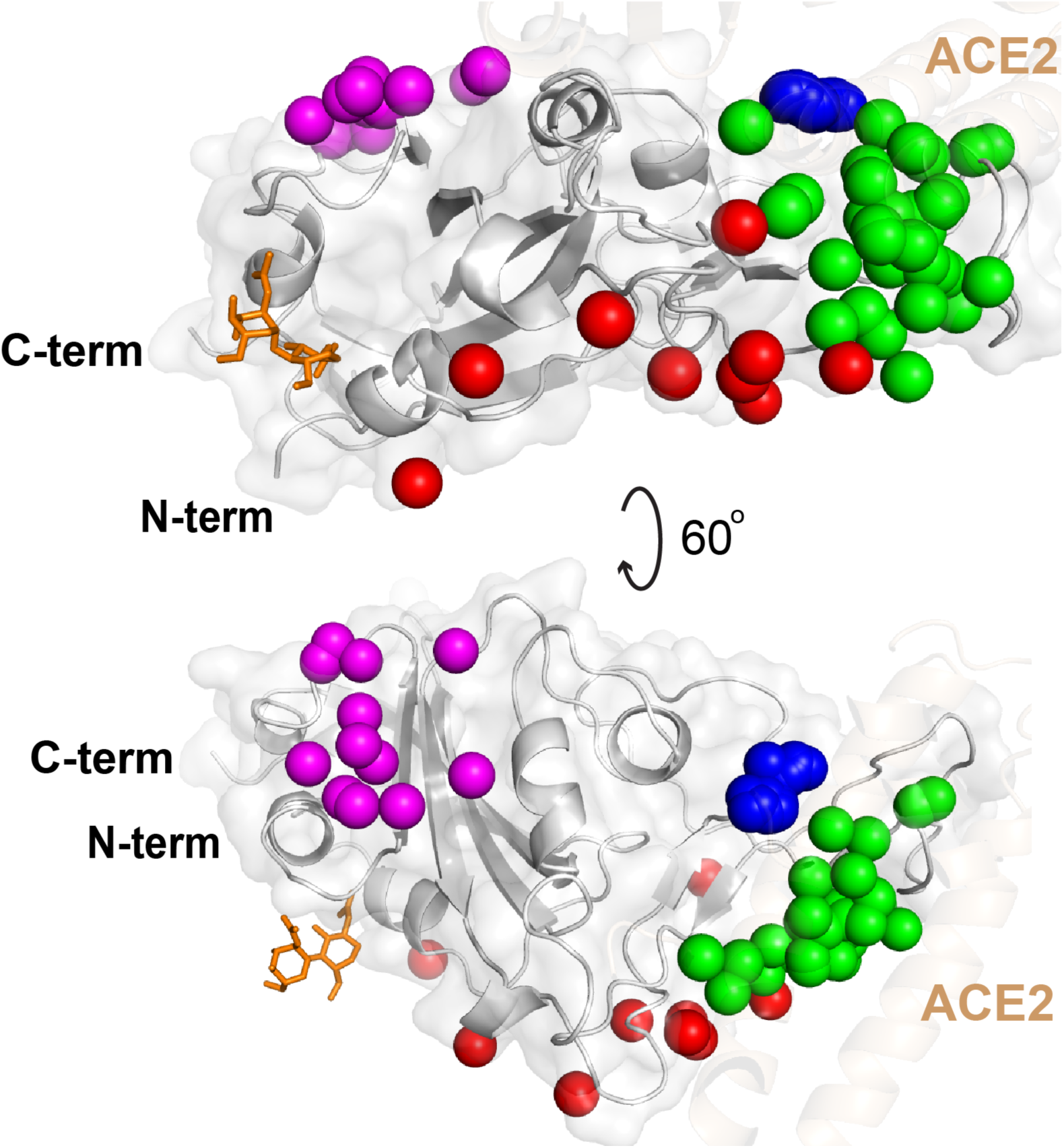
Distribution of antibody clusters on the receptor binding domain. Each antibody is represented as a sphere at the paratope center (centroid of all non-hydrogen atoms within 5Å of the RBD), and colored by contact-based antibody cluster (1: blue, 2: green, 3: red, 4: magenta). A representative RBD structure (from PDB code 7KN5) is shown in gray, and the N-glycan at residue N343 from that structure is shown as orange sticks. For reference, the superposed RBD-bound ACE2 structure (PDB code 6LZG) is shown as tan cartoon.

The contact-based clusters in **Figure 2** highlight several notable features within and between sets of RBD-targeting antibodies. Cluster 1 antibodies all neutralize SARS-CoV-2, block ACE2 binding, can only bind the spike in its open conformation, and have relatively high RBD interface buried surface area (BSA). Cluster 2 contains antibodies that can bind the closed spike, some of which can engage multiple RBDs in that context, and all are predicted or confirmed to block ACE2 binding. Cluster 3 is dominated by antibodies that can bind the closed spike, and most are predicted block ACE2 binding through steric hindrance and/or binding site overlap. In Cluster 4, which is mapped closer to the N- and C-termini and the hinge that connects the RBD to the spike (**Figure 3**), multiple antibodies are confirmed to be cross-neutralizing between SARS-CoV-2 and SARS-CoV-1 (Liu et al., 2020; Lv et al., 2020; Piccoli et al., 2020), and no antibodies are predicted to recognize spike in the RBD-closed conformation. The mapped antibody footprints show varying degrees of overlap with ACE2 binding site residues (gray bars at top of **Figure 2**) among the clusters. Residues highlighted in **Figure 2** that are associated with viral variants of concern (E484, K417, N501) show that Cluster 2 is primarily associated with E484 engagement, while Cluster 1 is associated with engagement of K417 and N501. Antibodies in Clusters 3 and 4 exhibit few or no contacts with those residues, suggesting that they are less susceptible to binding disruption and viral resistance due to variability at those sites.

### Binding energetic features and hotspots

To provide a more detailed and comprehensive view of key residues and energetic features underlying antibody-RBD recognition, all interface structures were analyzed for hydrogen bonds with RBD residues (**Figure 4**) and energetically important RBD residues based on computational alanine scanning (**Figure 5**). Hydrogen bonding patterns in RBD-targeting antibodies (**Figure 4**) showed clear preferences for hydrogen bond RBD residue interactions among Cluster 1 antibodies, with frequently observed interactions with residues R403, K417, D420, Y421, N487, and Y505. Many Cluster 2 antibodies exhibit hydrogen bond interactions with residue E484 and/or Q493, whereas antibodies from Clusters 3 and 4 have limited shared RBD residues involved in hydrogen bond interactions.

**Figure 4.**
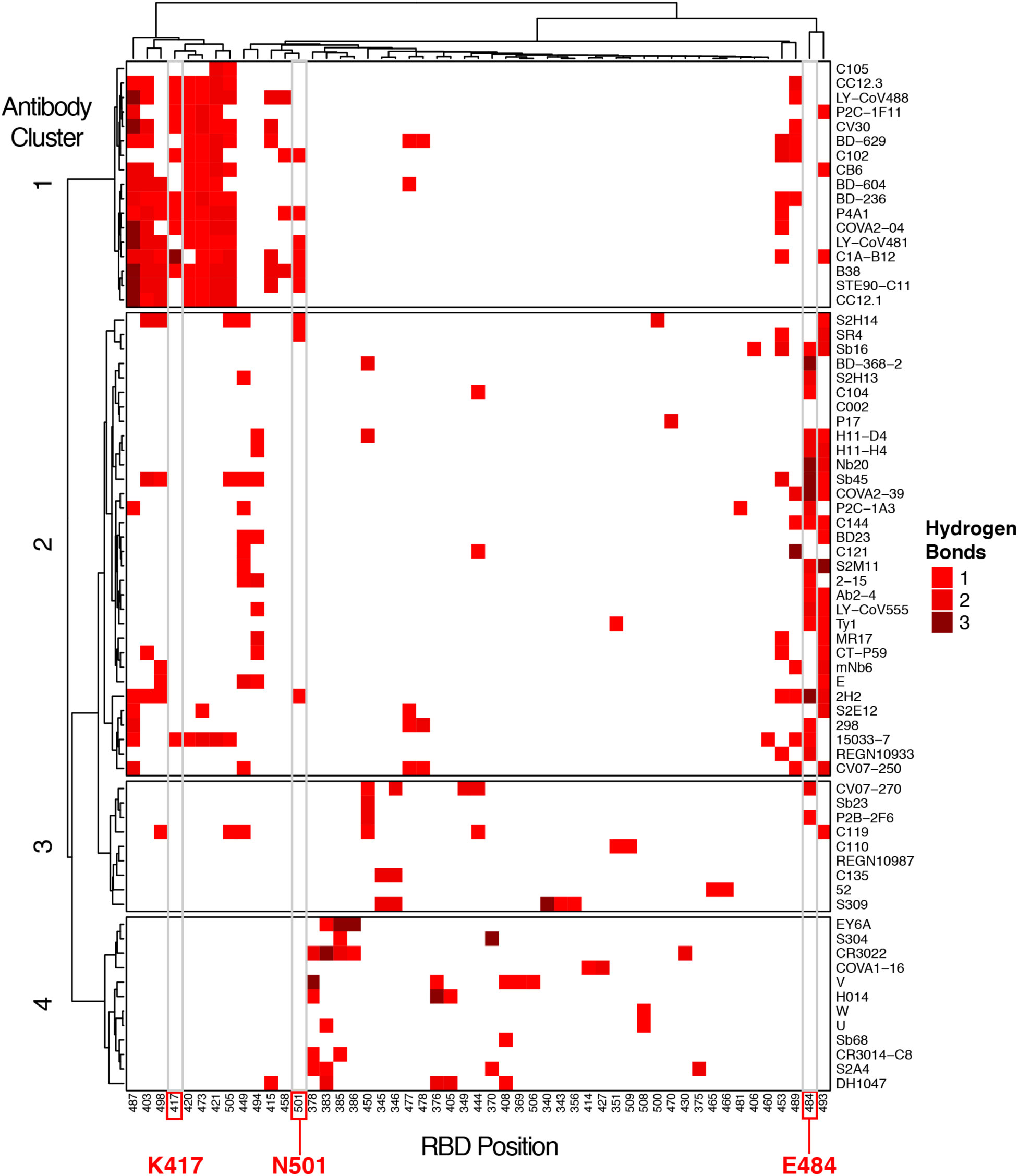
RBD hydrogen bond contacts of SARS-CoV-2 antibodies. Hydrogen bonds to RBD residue side chains were calculated for all antibody-RBD complexes using the hbplus program (McDonald and Thornton, 1994). Each hydrogen bond contact is colored by number of hydrogen bond interactions, as indicated on the key, and RBD positions are ordered by hierarchical clustering based on hydrogen bond profile similarities, with corresponding dendrogram shown at top. Antibodies (rows) are ordered and clustered as in Figure 2, based on the RBD contact profile similarities.

**Figure 5.**
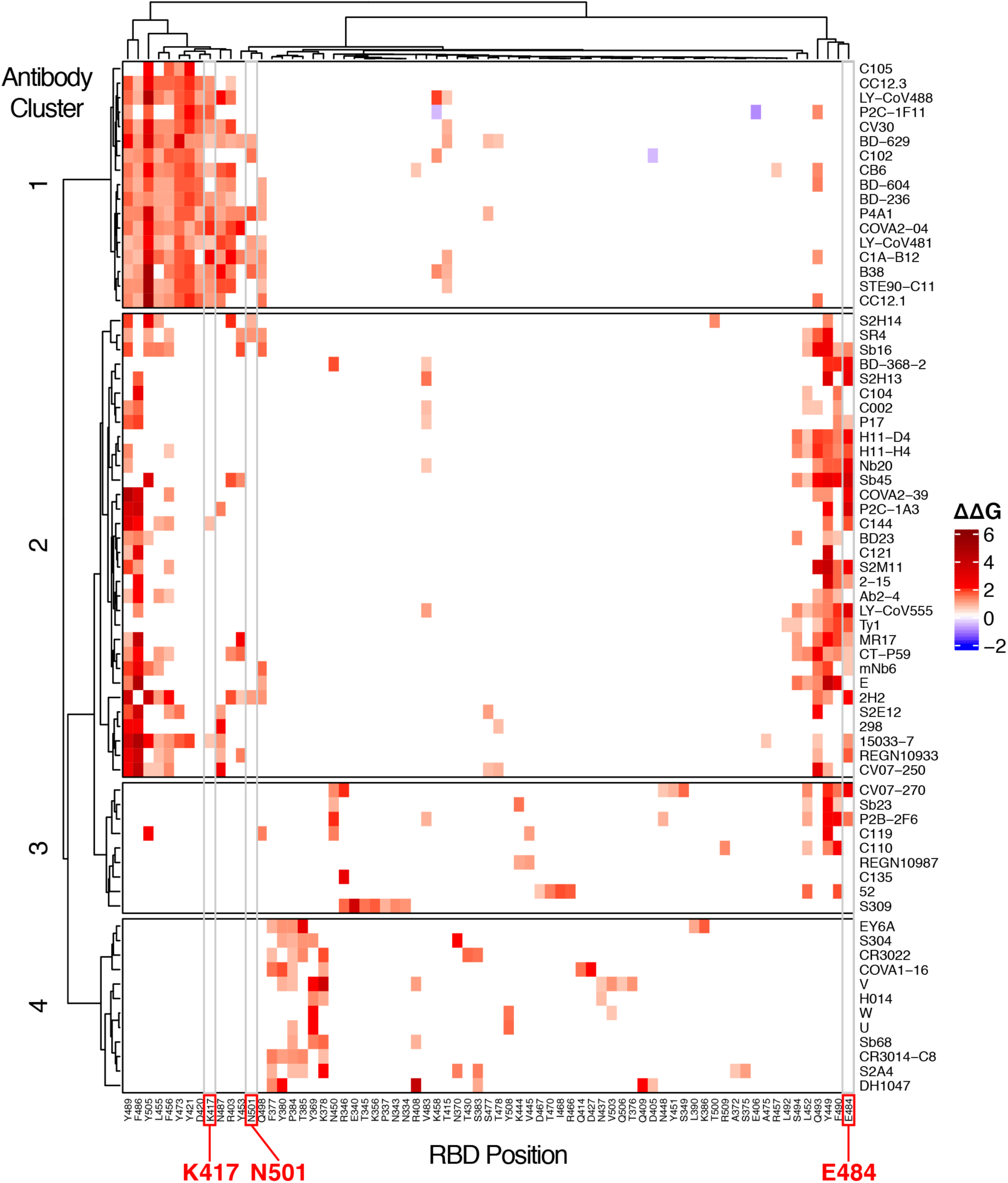
Computational mapping of SARS-CoV-2 RBD hotspot residues. Computational alanine scanning of RBD residues in antibody-RBD interfaces was performed using Rosetta (Kortemme et al., 2004), to generate binding energy change (ΔΔG) values for alanine substitutions at each RBD position based on modeling of residue substitutions and scoring using an energy-based function. ΔΔG values are in Rosetta Energy Units (REU) which are comparable to energies in kcal/mol. Alanine residues in the native complex were mutated to glycine for ΔΔG calculations, and glycine RBD residues were omitted from the analysis. In order to highlight substantial predicted binding energy changes, only ΔΔGs with absolute values > 0.5 REU are represented. RBD residues are ordered by hierarchical clustering based on ΔΔG profile similarities, with corresponding dendrogram shown at top. Antibodies (rows) are ordered and clustered as in Figure 2, based on the RBD contact profile similarities.

To map key RBD sites and energetic hotspots in the set of antibody-RBD interfaces, we performed computational alanine scanning (**Figure 5**) using a predictive protocol in Rosetta (Kortemme et al., 2004). The protocol used for this analysis was selected based on predictive performance from benchmarking of nine computational methods using approximately 350 experimentally determined alanine mutant ΔΔG values for antibody-antigen interfaces (**Table S2**). While many energetically important residues identified by this analysis are reflective of the key residues identified by hydrogen bond analysis, including residues N487 and E484 (Cluster 1) and E484 (Cluster 2), numerous hydrophobic RBD residues were additionally identified as important for binding within antibody clusters. These residues include Y505 (Cluster 1), F486 and Y489 (Clusters 1 and 2), and Y449 and F490 (Clusters 2 and 3). As with the analysis of RBD residue contacts, analysis of hydrogen bonds and computational alanine scanning support the overall importance of N417 and Y501 for Cluster 1 antibodies, and E484 for Cluster 2 antibodies.

### Epitope conservation and targeting of escape variants

To assess the degree to which antibodies and antibody classes to target sites that are conserved among sarbecoviruses, the fraction of RBD epitope residues conserved between SARS-CoV-2 and SARS-CoV-1 was calculated for each antibody-RBD interface (**Figure 6**). Clusters 1-3 exhibit limited conservation (approximately 50% or lower conserved antibody contact residues), with the exception of S309, which shows over 80% epitope residue conservation; this result is in accordance with the observed cross-neutralizing capability for that antibody (Pinto et al., 2020). In contrast with the other antibody clusters, antibodies in Cluster 4, which includes three confirmed cross-neutralizing antibodies (**Figure 2**), exhibit markedly higher epitope conservation, with all values 78% or higher. This highlights the potential importance of this conserved site, which is not accessible in the closed spike structure, in targeting of and immunity to emerging sarbecoviruses.

**Figure 6.**
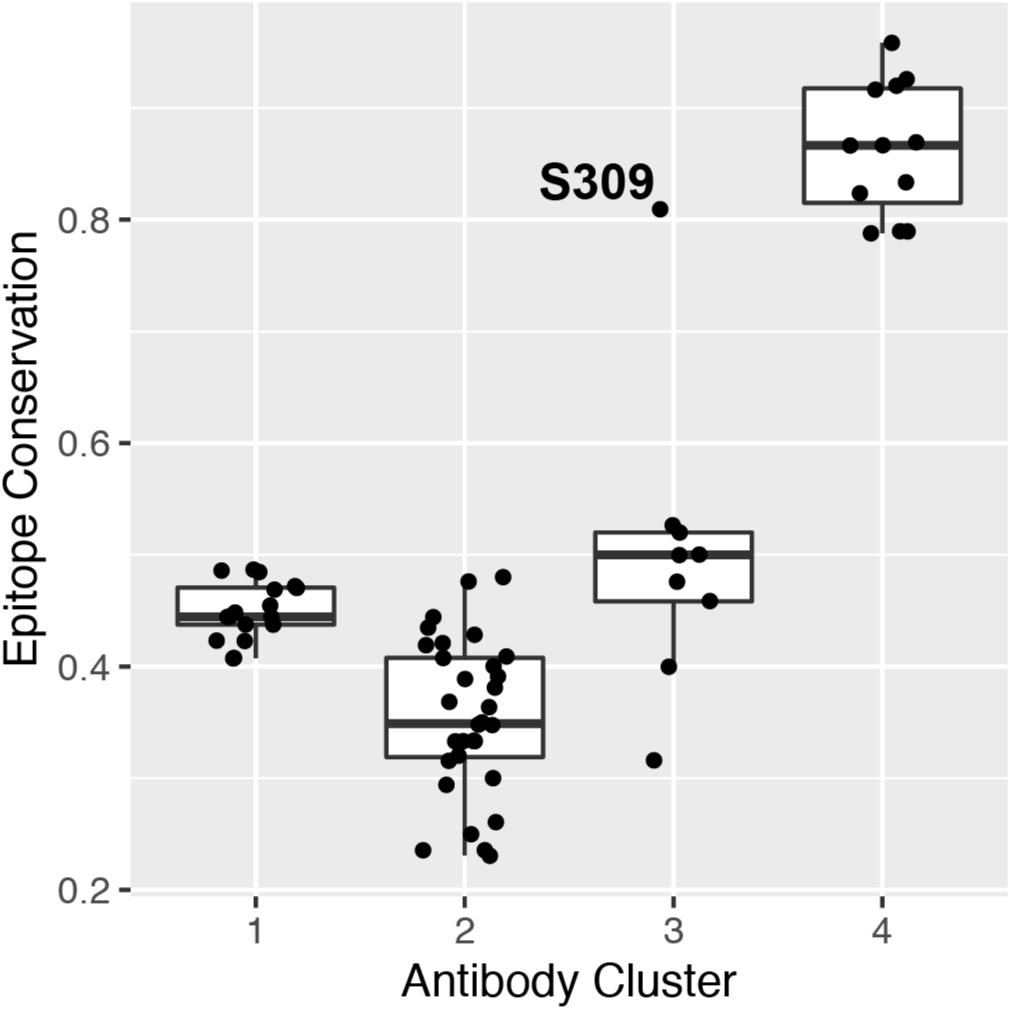
Epitope residue conservation in SARS-CoV-1 by antibody cluster. Epitope conservation, defined as the fraction of RBD epitope residues (< 5 Å distance to antibody) conserved between SARS-COV-1 and SARS-COV-2, was calculated for 70 antibody-RBD complex structures, and conservation values are shown as a boxplot grouped by antibody clusters, with all conservation values shown as points. The outlier point for Cluster 3 (S304 antibody) is labeled, and the total numbers of points are 17 (Cluster 1), 32 (Cluster 2), 9 (Cluster 3), and 12 (Cluster 4).

To directly assess the effects of RBD mutations present in recently described SARS-CoV-2 variants of concern, we performed computational mutagenesis to gauge whether antibody binding affinities are predicted to be disrupted by individual RBD substitutions. For these simulations, we utilized the same protocol that was used for computational alanine scanning; we found this method to have similar predictive performance for point residue substitutions to all residue types in comparison with performance for alanine-only substitutions (Pearson Correlation Coefficient (PCC) with experimental ΔΔGs of 0.5 for all residues, versus 0.53 for alanine-only; **Table S2**). RBD substitutions K417N, K417T, E484K, and N501Y were modeled in all interfaces and assessed for antibody ΔΔGs; these substitutions are collectively represented in variants B.1.1.7 (N501Y), B.1.351 (501Y.V2; K417N, E484K, N501Y), P1 (484K.V2; K417T, E484K, N501Y), B.1.525 (E484K), and a recently reported variant of concern, B.1.526 (E484K) (Annavajhala et al., 2021). Comparison of predicted ΔΔGs (**Figure 7**) shows that K417N, K417T, and to a lesser extent N501Y, are predicted to predominantly affect antibodies in Cluster 1, whereas disruptive effects of E484K are primarily observed for antibody Cluster 2, with the exception of two antibodies with predicted ΔΔG values of over 1 Rosetta Energy Unit (REU), in Cluster 3. In contrast, antibodies in Cluster 3 and 4 exhibit little overall effects from those variant RBD substitutions. We also tested predicted binding effects using a different modeling tool (FoldX), which uses a distinct modeling and scoring protocol from Rosetta, and found similar trends among antibody classes for the effects of the variants (**Figure S4**). Finally, two additional RBD substitutions (L452R, S477N) from other described SARS-CoV-2 variants that have unclear associations with transmissibility or antibody resistance (e.g. B.1.429+B.1.427, which contains L452R) were tested for effects in the antibody-RBD interfaces; neither was predicted to have a pronounced effect on recognition for any of the antibody clusters (**Figure S5**).

**Figure 7.**
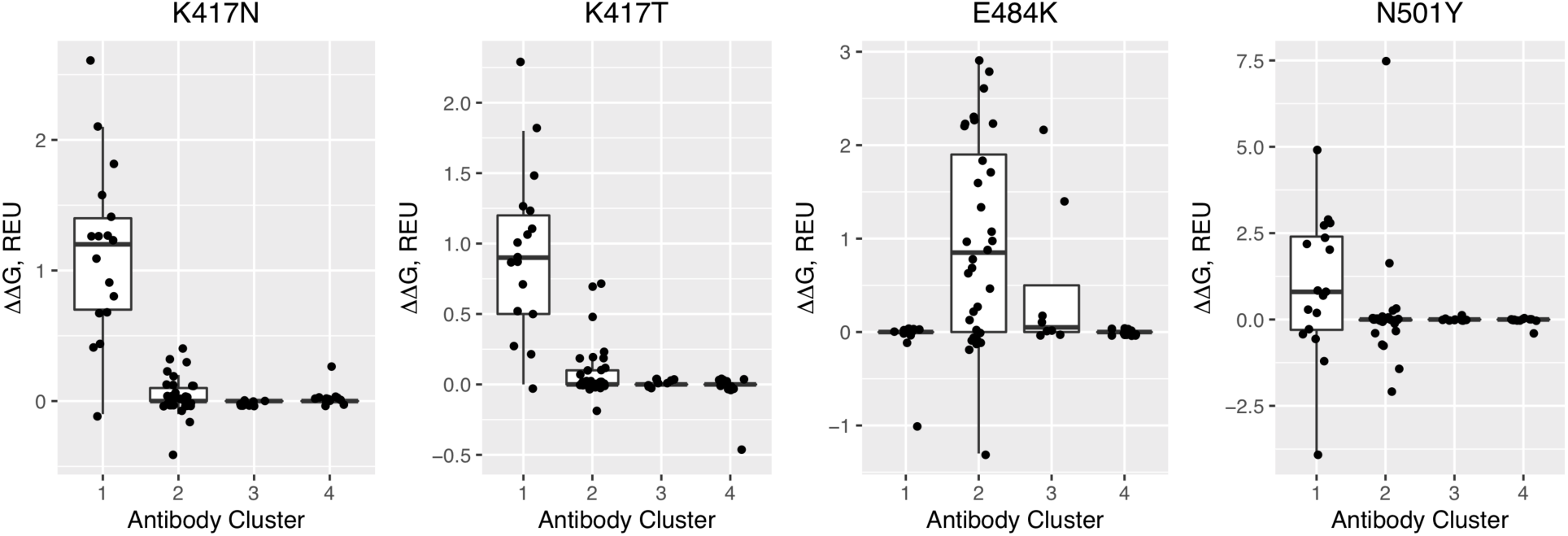
Profiling antibody binding disruption of RBD substitutions from circulating SARS- CoV-2 variants. Computational mutagenesis in Rosetta (Kortemme et al., 2004) was used to predict binding affinity effects (ΔΔGs) of RBD variant substitutions K417N, K417T, E484K, and N501Y for 70 antibodies that target the RBD. ΔΔG values are in Rosetta Energy Units (REU), which are comparable to energies in kcal/mol, and shown as boxplots grouped by antibody clusters, with all ΔΔG values shown as points. The total numbers of points are 17 (Cluster 1), 32 (Cluster 2), 9 (Cluster 3), and 12 (Cluster 4).

## DISCUSSION

Utilizing a curated set of experimentally determined antibody-RBD complex structures, we have performed detailed mapping of antibody recognition determinants on the SARS-CoV-2 RBD, which were used to generate antibody clusters that exhibit distinct structural and energetic signatures. Notably, these clusters exhibited different destabilizing effects from RBD substitutions found in circulating variants, underscoring and expanding upon previous observations by others that indicate that specific groups of antibodies are affected by specific substitutions, including E484K (Barnes et al., 2020a). We found that Cluster 2 antibodies, which overlap with Class 2 antibodies reported by Barnes et al. (**Figure 2**), are susceptible to resistance from viruses with the E484K substitution, which include B.1.351, P1, B.1.525, and B.1.526, but not B.1.1.7, whereas other antibodies are not likely to be affected by that substitution. In contrast, substitutions at residues K417 and N501, which are found in several variants of concern, were primarily associated with binding disruption to Cluster 1 antibodies based on our computational mutagenesis. Given that the E484K substitution appears specifically associated with viral escape, as noted by others (Altmann et al., 2021) and supported by recent studies of monoclonal and polyclonal antibody neutralization of variant viruses and specific mutants (Wang et al., 2021b; Wu et al., 2021), our work highlights the relative importance of Cluster 2 antibodies in the neutralizing response against SARS-CoV-2.

Though effects from RBD substitutions on ACE2 recognition were not considered in this study, due to its focus on antibody recognition and mutational escape, others have reported computational (Chen et al., 2020; Laurini et al., 2020) and experimental (Starr et al., 2020) RBD substitutions associated with loss of, or improvement of, ACE2 binding. As ACE2 binding effects can impact viral infectivity and fitness, a prospective combination of datasets from our study and a profile of ACE2 binding effects can provide a more comprehensive view of the landscape of viral fitness and immune escape. Such integrative work could identify SARS-CoV- 2 RBD variants with functional implications through computational structural analysis which are not yet identified in circulating variants, and can be prioritized for experimental characterization, and potentially with targeted therapies and updated vaccines, if they do appear. Additionally, new viral variant sequences can be rapidly assessed for possible mutational escape using our computational analysis pipeline.

This study is distinguished from other recently described structure-based (Barnes et al., 2020a) and binding competition-based (Dejnirattisai et al., 2021; Piccoli et al., 2020) reports to compare and classify antibodies, as we directly assessed detailed antibody binding footprints on the RBD with structural analysis to generate the identified clusters. The unsupervised clustering used here corroborated and expanded upon previously identified classes (Barnes et al., 2020a), though several distinctions in classifications were also observed in our analysis. To provide an updated reference to the community, we report these clusters on our CoV3D site of coronavirus protein structures (Gowthaman et al., 2021) (https://cov3d.ibbr.umd.edu/antibody_classification), which includes the 70 complexes reported in this study as well as newly reported complexes. We also provide a prototype interface on the CoV3D site for researchers to input new experimentally determined structures or models of antibody-RBD or protein-RBD complexes to characterize the binding footprint and provide the contact-based cluster.

New datasets reporting large-scale experimental mapping of antibody binding determinants can expand upon our analysis and provide additional insights. While in this work we report systematic computational alanine scanning to identify key energetic determinants of a large set of monoclonal antibodies that target the SARS-CoV-2, other studies have reported experimental global alanine scanning of antibody interactions with viral glycoproteins, such as hepatitis C virus E1E2, to map binding determinants (Colbert et al., 2019; Gopal et al., 2017; Keck et al., 2019; Pierce et al., 2016). These datasets were used to cluster antibodies and E1E2 positions by binding profiles in several of those studies (Colbert et al., 2019; Keck et al., 2019; Pierce et al., 2016). Though based on deep mutational scanning rather than direct measurement of binding affinities, recent studies provide information on the impact of RBD substitutions to alanine and other residues on recognition by sets of monoclonal antibodies (Greaney et al., 2021b; Starr et al., 2021), and such data could be inspected with respect to residue and mutation-level impact on antibody clusters identified here.

Certain elements of our analysis of antibody binding determinants can be expanded in future studies. Some omissions from the calculation of antibody contacts and energetic determinants on the RBD include lack of inclusion of certain non-protein atoms, such as water molecules and N-glycans, and lack of explicit calculation of adjacent RBD contacts, outside of noting likely closed spike cross-protomer binding, which was the case for a small fraction of antibodies considered here. Water molecules, which could mediate hydrogen bonds between antibody and RBD, were not included here, to avoid bias due to varying experimental structural resolutions which in many cases could not resolve water molecules, necessitating modeling of explicit water molecules which would lead to additional uncertainties in subsequent calculations (Lensink et al., 2014). Likewise, the N-glycans of the RBD, specifically the glycan at residue N343, has varying occupancies in experimentally determined structures. Though this glycan is contacted by the S309 antibody (Pinto et al., 2020), such antibody-RBD glycan contacts appear to be rare, at least for structurally characterized neutralizing antibodies, of which most compete with ACE2 binding and thus target regions sites that are not proximal to that N-glycan. One potential avenue for expansion of this analysis includes antibodies that target other regions of the spike glycoprotein, specifically the N-terminal domain (NTD). While the current set of experimentally determined SARS-CoV-2 antibody-NTD complex structures (March, 2021) is currently limited to six antibodies (4A8, FC05, DH1050.1, 2-51, COVOX-159, DH1052) (Cerutti et al., 2021; Chi et al., 2020; Dejnirattisai et al., 2021; Li et al., 2021; Wang et al., 2021a), recent structural and antigenic mapping studies of antibody recognition of this domain (Cerutti et al., 2021; McCallum et al.) indicate that a focused computational analysis would be useful, particularly as more structures of antibodies targeting this domain are reported. We currently represent this set in the “non-RBD” antibody class on the CoV3D site.

In addition to providing a view of the detailed landscape of antibody-RBD recognition determinants and key sites, our results indicate that certain sets of antibodies are associated with limited likely viral resistance from circulating variants (Clusters 3 and 4) as well as higher epitope sequence conservation (Cluster 4). Many of the antibodies in these sets have been experimentally confirmed to neutralize SARS-CoV-1 or cross-neutralize SARS-CoV-1 and SARS-CoV-2. Prospective structure-based antigen design studies could potentially focus the antibody response to the corresponding epitopes of the SARS-CoV-2 RBD, versus the epitopes collectively targeted by antibodies in Clusters 1 and 2. As binding of Cluster 4 antibodies is prevented in the context of the closed-RBD spike conformation, open spike antigen designs or RBD-only antigens would facilitate elicitation of these antibodies. Several recent studies have reported success using RBD displayed on self-assembling nanoparticles (Cohen et al., 2021; Walls et al., 2020a; Zhang et al., 2020), and structure-guided RBD optimization in the context of such a platform could lead to improved elicitation of desired antibody profiles. Integrating computational structural analysis and design with experimental characterization is a promising avenue toward effective combatting of SARS-CoV-2 variants and future emerging viruses.

## MATERIALS AND METHODS

### Structure assembly and curation

Structures of antibody-RBD complexes were downloaded from the CoV3D database (Gowthaman et al., 2021), which identifies and antibody-RBD structures in the Protein Data Bank (Rose et al., 2011) on a weekly basis through sequence similarity to coronavirus reference protein sequences in conjunction with identification and annotation of antibody chains. The set of antibody-RBD structures (downloaded in February 2021) was filtered for antibody nonredundancy based on antibody name and sequence identity, as well as resolution (< 4.0 Å). In cases of an antibody present in multiple antibody-RBD complex structures, the structure with highest resolution was selected for analysis. To permit consistency among antibody-RBD complex structures, and to facilitate calculations, antibodies were truncated to include variable domains, and full spike glycoproteins were truncated to include only RBD residues (residues 330-530) of the sole or major target of the antibody. To provide uniform input structures for atomic contact and other calculations, non-amino acid HETATMs were removed prior to structural analysis, and to resolve double occupancies and add missing side chain atoms, structures were pre-processed by the “score” application in Rosetta version 3.12 (Leman et al., 2020). Two complexes with missing side chain atoms in the experimental PDB coordinates were processed using the FastRelax protocol in Rosetta (Khatib et al., 2011), to perform constrained local minimization and to resolve unfavorable energies due to clashes from rebuilt side chains. These complexes Parameter flags used in FastRelax (“relax” executable in Rosetta 3.12) are:

-relax:constrain_relax_to_start_coords

-relax:coord_constrain_sidechains

-relax:ramp_constraints false

-ex1

-ex2aro

-no_optH false

-flip_HNQ

-renumber_pdb F

-nstruct 1

Antibody-RBD structures were aligned into a common reference frame through superposition of RBD coordinates using least-squares fitting in PyMOL (Schrodinger, Inc.). This set of pre- processed and aligned structures is available through the CoV3D site (Gowthaman et al., 2021), at: https://cov3d.ibbr.umd.edu/download (“Nonredundant RBD-antibody complex structures” link).

Information regarding neutralization of SARS-CoV-2 and SARS-CoV-1 was obtained from the CoV-AbDab site (Raybould et al., 2020), as well as references from the literature for certain antibodies, where noted in **Table S1**.

### Computational structural analysis

RMSD values between antibody heavy chain or nanobody orientations were determined by superposition of one antibody variable domain onto another using the FAST structure alignment tool (Zhu and Weng, 2005), and calculation of backbone RMSD between superposed and non-superposed variable domain (in the context of a common RBD reference frame, as noted above). Interface contacts are defined as inter-atomic distance between non-hydrogen atoms of less than 5 Å, and antibody-RBD residue contact maps were generated based on the total number of antibody atom contacts with each RBD residue. Hierarchical clustering of antibody RMSDs was performed in R version 4.0.3 (www.r-project.org) with the distance matrix of RMSDs as input, and Ward’s minimum variance method (“ward.D2” method in hclust). Hierarchical clustering of antibodies and RBD positions based on contact data was performed in R, using Manhattan distance to compute differences in contact profiles between antibodies or RBD positions, and Ward’s minimum variance method for clustering. Hierarchical clustering of RBD positions based on hydrogen bond or calculated ΔΔG values, for the respective heatmap figures, was likewise performed in R, using Manhattan distances and Ward’s clustering algorithm. RBD residue dimension reduction for representation in heatmap (**Figure 2**) was performed by selecting exemplar residues from 100 hierarchical clusters, which removed residues with highly similar contact profiles with respect to those shown in the heatmap. Generally, the omitted residues had low numbers of total antibody contacts, as Manhattan distances, based on total atomic contacts with each antibody, were used for contact-based distance calculations between RBD residues. The pvclust method (Suzuki and Shimodaira, 2006), as implemented in R, was used to calculate bootstrap confidence of contact-based hierarchical clusters of antibodies, using 20,000 bootstrap replicates. Principal component analysis of antibody-RBD contact profile data was performed with the scikit-learn Python module.

Buried surface areas (BSAs) were calculated using the naccess program (v. 2.1.1) (Hubbard and Thornton, 1993), subtracting the solvent accessible surface area of the antibody-RBD complex structure from the total solvent accessible surface area of the separate antibody and RBD structures, dividing by two to avoid double-counting interface area and to make BSA values commensurate with those from other tools including PISA (http://www.ebi.ac.uk/pdbe/prot_int/pistart.html). Antibody-RBD interface hydrogen bonds were calculated using the hbplus program (v. 3.15) (McDonald and Thornton, 1994), with default parameters.

Structure-based calculations of antibody blocking of ACE2 binding to RBD were calculated using the ACE2-RBD complex structure (PDB code 6LZG) (Wang et al., 2020). After superposition of ACE2-RBD and antibody-RBD complexes by RBD, the number of inter-atomic clashes, defined as non-hydrogen atom pairs with distances < 2.5 Å, was calculated between ACE2 and each antibody structure. Antibodies with > 20 atomic clashes with ACE2 were classified as likely to block ACE2 binding. Structure-based calculations of antibody binding to the closed spike structure were performed using the SARS-CoV-2 closed spike structure reported by Walls et al. (PDB code 6VXX) (Walls et al., 2020b). Antibodies with < 100 atomic clashes with spike atoms outside of the target RBD structure and chain after superposition of the antibody-RBD complex onto the 6VXX structure were classified as predicted to bind the closed spike. Clash thresholds were selected based on agreement with structures and experimental data regarding ACE2 blocking and closed spike binding, when available. Four antibodies that engaged the closed spike and exhibited cross-protomer binding, as confirmed by inspection of antibody-spike complex structures (S2M11, C144, mNb6, LY-CoV555; PDB codes 7K43, 7K90, 7KKL, 7L3N) (Barnes et al., 2020a; Jones et al., 2020; Schoof et al., 2020; Tortorici et al., 2020), were annotated accordingly in the contact heatmap.

### Computational mutagenesis

Computational modeling and prediction of antibody binding energy changes (ΔΔGs) for alanine and other residue point substitutions was performed using Rosetta version 2.3 (Kortemme et al., 2004), Rosetta version 3.12 (Leman et al., 2020), and FoldX version 4 (Schymkowitz et al., 2005). Benchmarking of computational alanine scanning predictive performance was performed using a subset of the AB-Bind dataset (Sirin et al., 2016) that contains alanine point substitutions with quantified experimental ΔΔG measurements and known wild-type complex structures (347 mutants and ΔΔG values). A larger set with all point substitutions (including non-alanine substitutions) was also tested (531 mutants and ΔΔG values). Pearson correlation coefficients (PCC) between measured and predicted ΔΔG values, and receiver operating characteristic area under the curve (AUC) values for prediction of hotspot residues (measured ΔΔG for alanine residue substitution > 1 kcal/mol), were calculated using scipy and scikit-learn (sklearn) Python libraries, respectively.

Prior to running ΔΔG calculations in Rosetta, antibody-RBD complex structures were pre- processed using Rosetta’s FastRelax protocol (Khatib et al., 2011), using the flags noted above, to perform backbone and side chain constrained minimization to resolve unfavorable energies and anomalies that would bias energetic calculations. Rosetta 2.3 ΔΔG calculations were performed using the “interface” protocol (Kortemme and Baker, 2002; Kortemme et al., 2004). An example command line is: rosetta.mactel -interface -intout pdb.ddgs.out -ignore_unrecognized_res -safety_check - skip_missing_residues -mutlist pdb.muts.txt -extrachi_cutoff 1 -ex1 -ex2 -ex3 -constant_seed - jran 12 -yap -s input.pdb

The input files specified on the command line denote the input PDB file (“input.pdb”) and the list of mutations (“pdb.muts.txt”). The default protocol only models the mutant residue for ΔΔG calculation (“Ros2.3_norepack” in **Table S2**), and additional flags were used on the command line to perform minimization of mutation-proximal side chains (“-min_interface -int_chi” flags; “Ros2.3_minint_chi” in **Table S2**), minimization of mutation-proximal side chains and backbone (“-min_interface -int_bb -int_chi” flags; “Ros2.3_minint_bb_chi” in **Table S2**), and rotamer-based packing of mutation-proximal side chains (“-repack” flag, “Ros2.3_repack” in **Table S2**).

Rosetta 3 ΔΔG calculations were performed with two available computational mutagenesis protocols. One Rosetta 3 computational alanine scanning protocol was downloaded from a public resource containing benchmarks and Rosetta tools (S et al., 2015), and represents a separate implementation of the Rosetta 2.3 mutagenesis protocol noted above (Kortemme and Baker, 2002; Kortemme et al., 2004). This protocol was recently used to predict TCR-peptide-MHC interface ΔΔG values (Wu et al., 2020). In addition to the default protocol that does not repack neighboring side chains (“Ros3_norepack” in **Table S2**), we also tested this protocol with repacking of neighboring side chains (“Ros3_repack” in **Table S2**).

An example command line for running this protocol is:

rosetta_scripts.static.linuxgccrelease -s input.pdb -parser:protocol alascan.xml -parser:view - inout:dbms:mode sqlite3 -inout:dbms:database_name rosetta_output.db3 -no_optH true - parser:script_vars pathtoresfile=input.resfile chainstomove=1,2 -ignore_zero_occupancy false

We additionally tested the alanine scan using the flex ddG protocol, which was developed recently in Rosetta 3 (Barlow et al., 2018). This protocol uses the backrub algorithm (Smith and Kortemme, 2008) to sample protein backbone conformations at the interface, and the average ΔΔG values are calculated over a number of models. We tested two sets of ΔΔG scores that are output by flex ddG, representing different scoring functions reported by the authors (Barlow et al., 2018); they are shown as “flex_ddG-fa_talaris2014” and “flex_ddG-fa_talaris2014-gam” in **Table S2**.

An example command line used for flex ddG calculations in this study is: rosetta_scripts.linuxgccrelease -s input.pdb -parser:protocol flexddg.xml -parser:script_vars chainstomove=1,2 mutate_resfile_relpath=input.resfile number_backrub_trials=35000 max_minimization_iter=5000 abs_score_convergence_thresh=1.0 backrub_trajectory_stride=7000 -restore_talaris_behavior -in:file:fullatom - ignore_unrecognized_res -ignore_zero_occupancy false -ex1 -ex2

For ΔΔG calculations in FoldX (Schymkowitz et al., 2005), complex structures were pre- processed using the FoldX RepairPDB protocol, and ΔΔG values were calculated using the FoldX PSSM protocol.

### Sequence conservation

Assessment of sequence conservation of SARS-CoV-2 RBD positions in the SARS-CoV-1 sequence was performed using SARS-CoV-2 (GenBank: QHD43416) and SARS-CoV-1 (GenBank: AAP13441) spike reference sequences aligned with BLAST (Altschul et al., 1990). The epitope residues of each antibody were defined as any SARS-CoV-2 residue within 5 Å of any antibody residue. An in-house Perl script was used to analyze SARS-CoV-2 antibody-antigen interfaces and calculate epitope conservation.

### Figures

Figures of structures were generated using PyMOL version 1.8 (Schrodinger, Inc.). Boxplots and dendrograms were generated using the ggplot2 (Wickham, 2016) and factoextra (Kassambara and Mundt, 2020) packages in R, and heatmaps were generated using the ComplexHeatmap package (Gu et al., 2016) in R.

## ACKNOWLEDGEMENTS

Computing resources from the University of Maryland Institute for Bioscience and Biotechnology Research High Performance Computing Cluster were used in this study. This work was supported by startup funding from the University of Maryland (B.G.P.) and NIH R01 GM126299 (B.G.P.).

## Notes

### Competing Interest Statement

The authors have declared no competing interest.

