## Supplemental Tables and Figures for "Structural and energetic profiling of SARS-CoV-2 antibody recognition and the impact of circulating variants"

**Table S1**. Antibody-spike and antibody-RBD complex structures analyzed in this study.

| Antibody name | PDB code | Type[1] | Species[2] | IGHV gene[2] | Neut[3] | Resolution (Å)[4] | Release Date[4] |
|---|---|---|---|---|---|---|---|
| Ab2-4 | 6XEY | ab | human | IGHV1-2 | Y | 3.25 | 7/21/20 |
| BD23 | 7BYR | ab | human | IGHV7-4-1 | Y | 3.84 | 6/9/20 |
| B38 | 7BZ5 | ab | human | IGHV3-66 | Y | 1.84 | 5/12/20 |
| BD-236 | 7CHB | ab | human | IGHV3-53 | Y | 2.4 | 9/15/20 |
| BD-368-2 | 7CHF | ab | human | IGHV3-23 | Y | 2.67 | 9/15/20 |
| BD-604 | 7CHF | ab | human | IGHV3-53 | Y | 2.67 | 9/15/20 |
| BD-629 | 7CH5 | ab | human | IGHV3-53 | Y | 2.7 | 9/15/20 |
| C105 | 6XCM | ab | human | IGHV3-53 | Y | 3.42 | 6/30/20 |
| CB6 | 7C01 | ab | human | IGHV3-66 | Y | 2.88 | 5/26/20 |
| CC12.1 | 6XC3 | ab | human | IGHV3-53 | Y | 2.7 | 7/7/20 |
| CC12.3 | 6XC4 | ab | human | IGHV3-53 | Y | 2.34 | 7/7/20 |
| COVA2-04 | 7JMO | ab | human | IGHV3-53 | Y | 2.36 | 8/25/20 |
| COVA2-39 | 7JMP | ab | human | IGHV3-53 | Y (Brouwer et al., 2020) | 1.71 | 8/25/20 |
| CR3022 | 6YLA | ab | human | IGHV5-51 | N | 2.42 | 4/14/20 |
| CV30 | 6XE1 | ab | human | IGHV3-53 | Y | 2.75 | 6/30/20 |
| EY6A | 6ZCZ | ab | human | IGHV3-30-3 | Y | 2.65 | 6/23/20 |
| H014 | 7CAI | ab | human | IGHV1-69-2 | Cross | 3.49 | 9/22/20 |
| H11-D4 | 6YZ5 | nano | llama | IGHV3-3 | Y | 1.8 | 6/2/20 |
| H11-H4 | 6ZH9 | nano | llama | IGHV3-3 | Y | 3.31 | 9/1/20 |
| MR17 | 7C8W | nano | alpaca | IGHV3S53 | Y | 2.77 | 6/23/20 |
| P2B-2F6 | 7BWJ | ab | human | IGHV4-38-2 | Y | 2.85 | 6/2/20 |
| REGN10933 | 6XDG | ab | human | IGHV3-11 | Y | 3.9 | 6/23/20 |
| REGN10987 | 6XDG | ab | human | IGHV3-30 | Y | 3.9 | 6/23/20 |
| S309 | 7JX3 | ab | human | IGHV1-18 | Cross | 2.65 | 10/14/20 |
| SR4 | 7C8V | nano | alpaca | IGHV3-3 | Y | 2.15 | 6/23/20 |
| Ty1 | 6ZXN | nano | alpaca | IGHV3-48 | Y | 2.93 | 9/22/20 |
| S2M11 | 7K43 | ab | human | IGHV1-58 | Y | 2.6 | 10/7/2020 |
| S2E12 | 7K4N | ab | human | IGHV1-2 | Y | 3.3 | 10/7/2020 |
| S2A4 | 7JVC | ab | human | IGHV3-7 | Y | 3.3 | 10/14/20 |
| S2H13 | 7JV6 | ab | human | IGHV3-7 | Y | 3 | 10/14/20 |
| COVA1-16 | 7JMW | ab | human | IGHV1-46 | Cross | 2.89 | 10/14/20 |
| CV07-250 | 6XKQ | ab | human | IGHV1-18 | Y | 2.55 | 10/14/20 |
| CV07-270 | 6XKP | ab | human | IGHV3-11 | Y | 2.72 | 10/14/20 |
| S304 | 7JX3 | ab | human | IGHV3-13 | Cross | 2.65 | 10/14/20 |
| S2H14 | 7JX3 | ab | human | IGHV3-15 | Y | 2.65 | 10/14/20 |
| C144 | 7K90 | ab | human | IGHV3-53 | Y | 3.24 | 10/21/20 |
| C135 | 7K8Z | ab | human | IGHV3-30 | Y | 3.5 | 10/21/20 |
| C121 | 7K8X | ab | human | IGHV1-2 | Y | 3.9 | 10/21/20 |

| | | | | | | | |
|---|---|---|---|---|---|---|---|
| C119 | 7K8W | ab | human | IGHV1-46 | Y | 3.6 | 10/21/20 |
| C110 | 7K8V | ab | human | IGHV5-51 | Y | 3.8 | 10/21/20 |
| C002 | 7K8T | ab | human | IGHV3-30 | Y | 3.4 | 10/21/20 |
| C102 | 7K8M | ab | human | IGHV3-53 | Y | 3.2 | 10/21/20 |
| Sb23 | 7A29 | nano | alpaca | IGHV3-3 | Y | 2.94 | 10/21/20 |
| C104 | 7K8U | ab | human | IGHV4-34 | Y | 3.8 | 10/21/20 |
| 298 | 7K9Z | ab | human | IGHV1-2 | Y | 2.95 | 10/28/20 |
| 52 | 7K9Z | ab | human | IGHV1-69 | Y | 2.95 | 10/28/20 |
| mNb6 | 7KKL | nano | alpaca | IGHV3S53 | Y | 2.85 | 11/11/20 |
| P2C-1A3 | 7CDJ | ab | human | IGHV3-11 | Y | 3.4 | 11/18/20 |
| P2C-1F11 | 7CDI | ab | human | IGHV3-66 | Y | 2.96 | 11/18/20 |
| P4A1 | 7CJF | ab | human | IGHV3-53 | Y | 2.11 | 11/11/2020 |
| C1A-B12 | 7KFV | ab | human | IGHV3-53 | Y | 2.1 | 12/2/2020 |
| Nb20 | 7JVB | nano | alpaca | IGHV3-3 | Y | 3.29 | 12/2/2020 |
| 2H2 | 7DK4 | ab | mouse | IGHV2-5-1 | Y | 3.8 | 12/2/2020 |
| P17 | 7CWN | ab | human | IGHV1-69-2 | Y | 3.2 | 12/16/2020 |
| STE90-C11 | 7B3O | ab | human | IGHV3-66 | Y | 2 | 12/16/2020 |
| CR3014-C8 | 7KZB | ab | human | IGHV3-72 | N | 2.83 | 2/3/2021 |
| Sb16 | 7KGK | nano | alpaca | IGHV3S53 | Y | 2.6 | 2/3/2021 |
| Sb45 | 7KGJ | nano | alpaca | IGHV3S53 | N | 2.3 | 2/3/2021 |
| DH1047 | 7LD1 | ab | human | IGHV1-46 | Y | 3.4 | 1/27/2021 |
| LY-CoV481 | 7KMI | ab | human | IGHV3-53 | Y | 1.73 | 1/27/2021 |
| LY-CoV488 | 7KMH | ab | human | IGHV3-53 | Y | 1.72 | 1/27/2021 |
| LY-CoV555 | 7KMG | ab | human | IGHV1-69 | Y | 2.16 | 1/27/2021 |
| W | 7KN7 | nano | alpaca | IGHV3-3 | Y | 2.73 | 1/20/2021 |
| V | 7KN6 | nano | alpaca | IGHV3S1 | Y | 2.55 | 1/20/2021 |
| E | 7KN5 | nano | alpaca | IGHV3-3 | Y | 1.87 | 1/20/2021 |
| U | 7KN5 | nano | alpaca | IGHV3-3 | Y | 1.87 | 1/20/2021 |
| CT-P59 | 7CM4 | ab | human | IGHV2-70 | Y (Kim et al., 2021) | 2.71 | 1/20/2021 |
| 2-15 | 7L5B | ab | human | IGHV1-2 | Y | 3.18 | 2/10/2021 |
| 15033-7 | 7KLH | ab | human | IGHV3-23 | Y | 3 | 2/10/2021 |
| Sb68 | 7KLW | nano | alpaca | IGHV3S53 | Y | 2.6 | 2/3/2021 |

[1]Antibody type. "ab": heavy-light chain antibody, "nano": nanobody/VHH.
[2]Species and IGHV gene name determined by ANARCI from antibody heavy chain or nanobody sequence.
[3]Measured SARS-CoV-2 neutralization, from CoV-AbDab (Raybould et al., 2020) or the literature, where specified by a reference. N: does not neutralize SARS-CoV-2; Y: neutralizes SARS-CoV-2; Cross: neutralizes SARS-CoV-2 and SARS-CoV-1.
[4]Resolution and release date of structure in the Protein Data Bank (PDB) (Rose et al., 2011).

**Table S2**. Performance of computational alanine scanning ΔΔG prediction for antibody-antigen interfaces.

| Method | Correlation[1] | AUC[2] |
|---|---|---|
| FoldX | 0.49 | 0.67 |
| Ros2.3_norepack | 0.52 | 0.71 |
| Ros2.3_minint_bb_chi | 0.50 | 0.70 |
| **Ros2.3_minint_chi** | **0.53** | **0.72** |
| Ros2.3_repack | 0.51 | 0.70 |
| Ros3_norepack | 0.44 | 0.68 |
| Ros3_repack | 0.45 | 0.69 |
| flex_ddG-fa_talaris2014 | 0.48 | 0.66 |
| flex_ddG-fa_talaris2014-gam | 0.52 | 0.69 |
| *Including non-alanine mutants* | | |
| Ros2.3_minint_chi[3] | 0.50 | 0.70 |

Computational predictions of binding affinity changes were computed for a subset of the AB-Bind dataset of measured antibody-antigen ΔΔG values (Sirin et al., 2016) with alanine point substitutions, available wild-type complex structures, and quantified ΔΔG measurements (347 total ΔΔGs). Rosetta version 2.3 (Ros2.3) (Kortemme et al., 2004), Rosetta version 3.12 (Ros3) (Leman et al., 2020), and FoldX (Schymkowitz et al., 2005) were used to predict ΔΔG values using various modeling and scoring protocols, as detailed in the Methods. Protocol selected for subsequent RBD alanine scanning based on performance comparison is shown in **bold**.

[1]Pearson correlation coefficient between predicted and experimentally determined ΔΔG values.

[2]ROC AUC value for prediction of hotspot (experimental ΔΔG > 1 kcal/mol) versus non-hotspot residues based on predicted ΔΔG values.

[3]Predictive performance for larger AB-Bind set that includes non-alanine point substitutions (531 mutants and ΔΔG values).

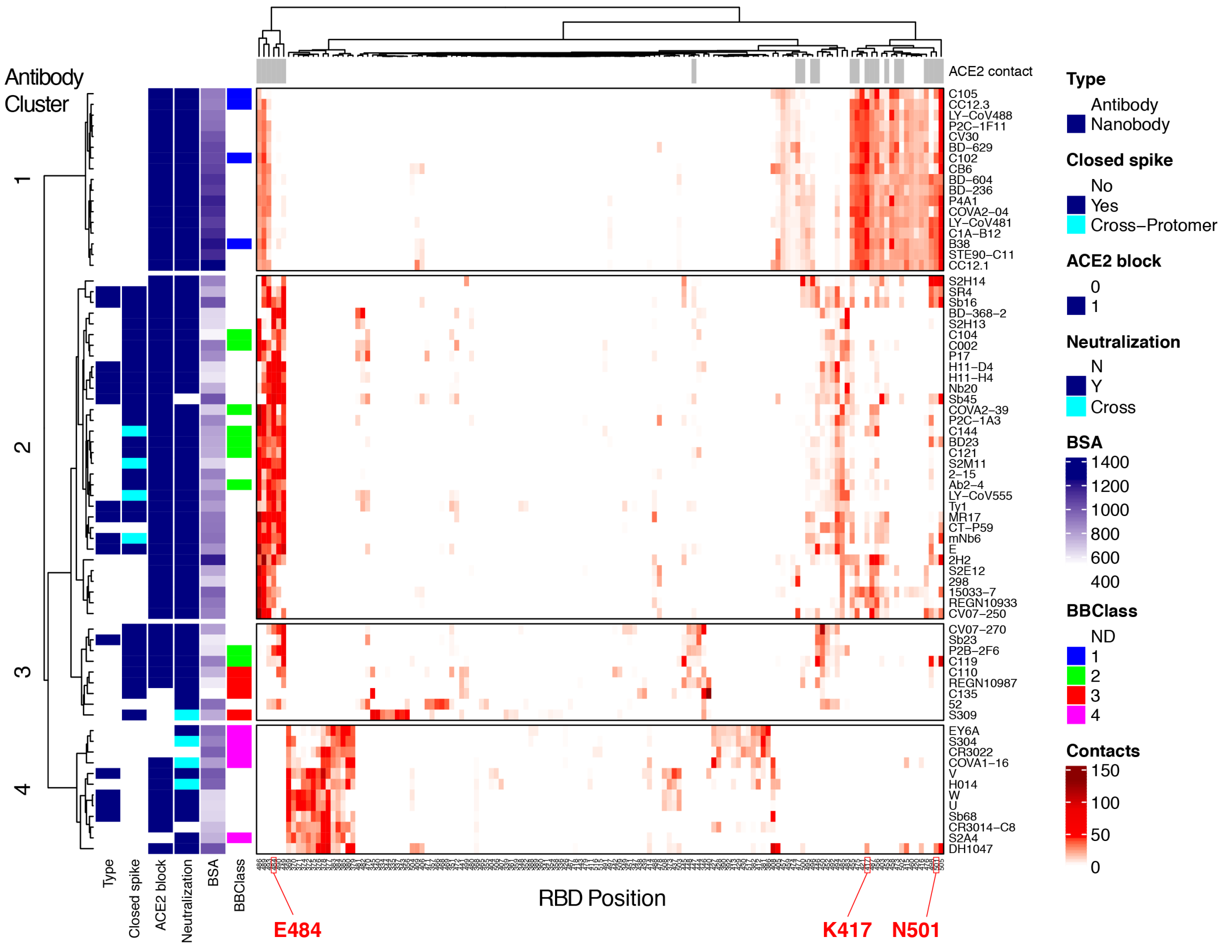

**Figure S1**. Heatmap of antibody-RBD contacts, representing the number of antibody atom contacts with each RBD residue and the full set of 139 contacted RBD positions. Labels and annotations are in accordance with the corresponding labels/annotations in **Figure 2**, and antibodies (rows) and RBD positions (columns) are ordered by hierarchical clustering in R. Antibodies in the heatmap are separated by the four major hierarchical clusters, which are labeled on left.

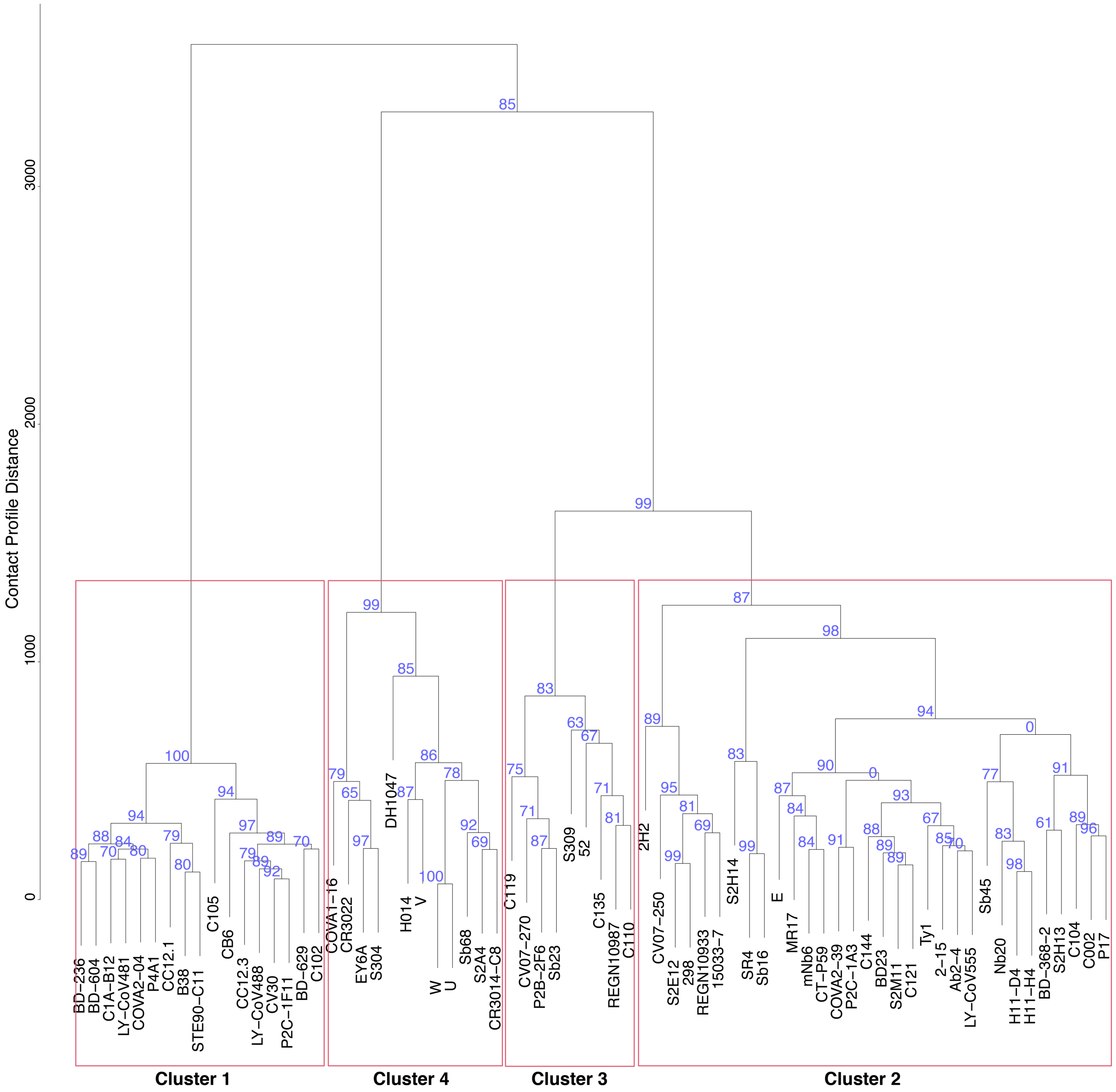

**Figure S2**. Antibody hierarchical clustering bootstrap confidence values, calculated by pvclust (Suzuki and Shimodaira, 2006). Multiscale bootstrap resampling was performed in pvclust in R, with the antibody-RBD contact data and 10,000 replicates. Values at each node denote the Approximately Unbiased (AU) bootstrap confidence, and red boxes delineate the four major clusters noted in this study, labeled accordingly.

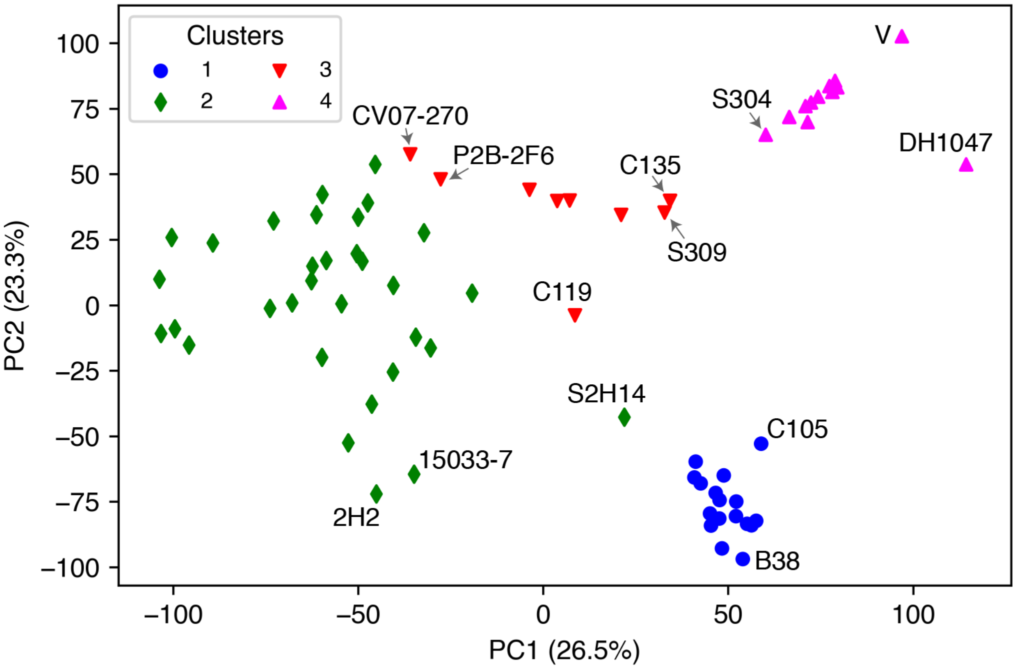

**Figure S3**. Principal component analysis of antibody-RBD residue footprint data. The x and y axes represent the first two principal components (PC1, PC2), with percentage of data variance represented by each principal component shown in parentheses. The 70 antibodies are shown as points, with colors and shapes representing Clusters 1-4, which were determined by hierarchical clustering analysis of antibody-RBD contact profiles. Selected points representing antibodies that are located on peripheries of cluster distributions are labeled by corresponding antibody names.

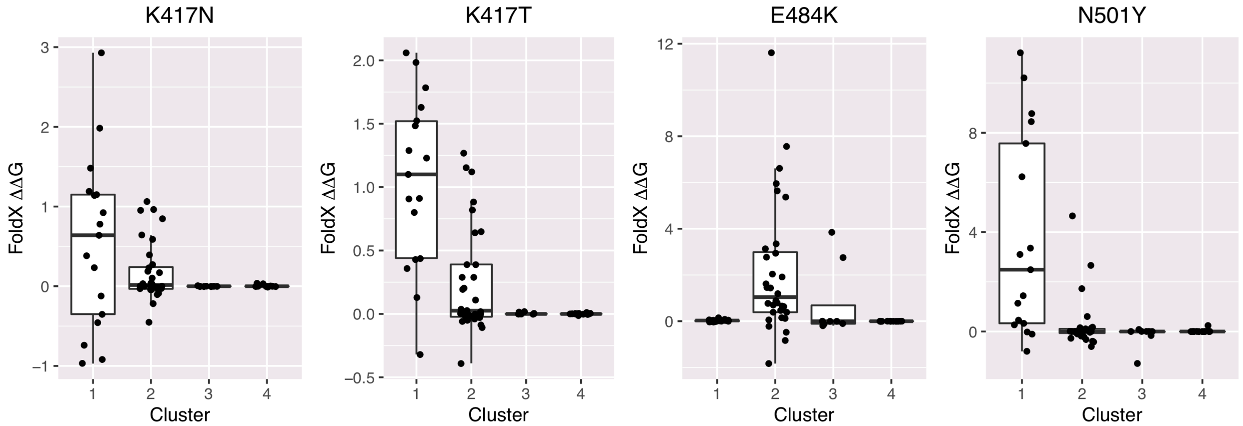

**Figure S4.** Computational assessment of antibody ΔΔG values for RBD variants using FoldX (Schymkowitz et al., 2005). FoldX was used to simulate and compute binding affinity changes (ΔΔGs) for RBD substitutions K417N, K417T, E484K, and N501Y individually in 70 antibody RBD interface structures. ΔΔG values for each RBD substitution are shown as a separate boxplot, with antibodies grouped by contact based cluster (1-4).

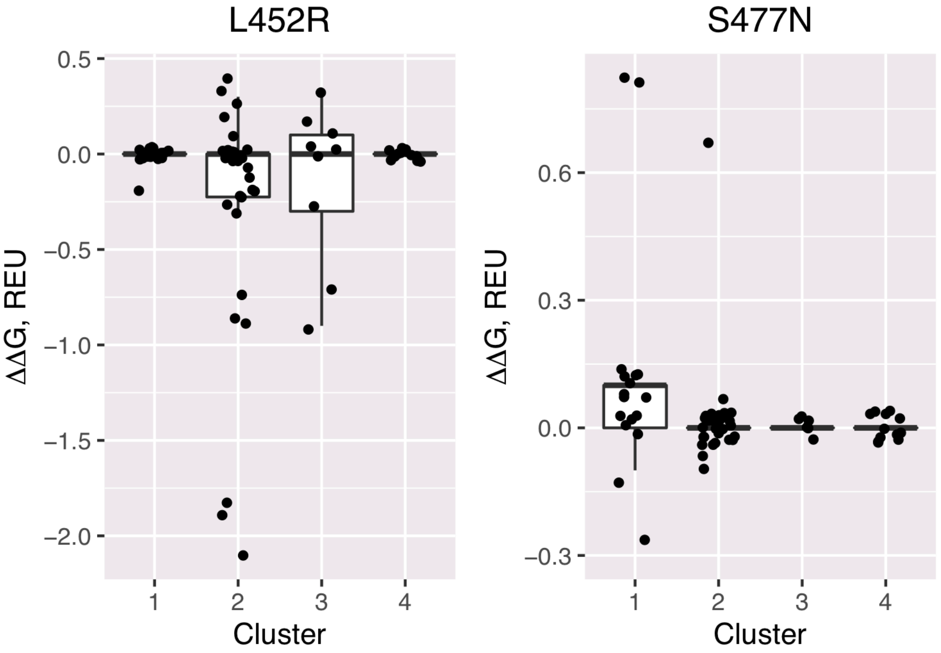

**Figure S5.** Calculated antibody binding affinity changes (ΔΔGs) for additional RBD variants using Rosetta. A mutagenesis protocol in Rosetta (Kortemme et al., 2004) was used to simulate and compute binding affinity changes (ΔΔGs) for RBD substitutions L452R and S477N individually in 70 antibody RBD interface structures. ΔΔG values for each RBD substitution are shown as a separate boxplot, with antibodies grouped by contact based cluster (1-4).
